# *Anopheles stephensi* bionomics and epidemiology in Ethiopia: A systematic review and meta-analysis with implications for urban malaria control

**DOI:** 10.64898/2026.04.15.718636

**Authors:** Teresa Beyena, Kidane Lelisa, Abdissa Deberssa, Lemma Regesa

## Abstract

**Background:** *Anopheles stephensi*, an invasive malaria vector originally endemic to South Asia, has rapidly expanded across East Africa. Its emerging malaria threats in urban Ethiopia threaten the elimination efforts, especially in areas once deemed low risk. A systematic review was conducted to synthesize the evidence regarding its bionomics and epidemiological impact, highlighting implications for urban malaria control strategies. This systematic review provides the first Ethiopia-specific quantitative evidence synthesis, addressing a vital knowledge gap necessary for guiding national malaria elimination programmes.

**Methods:** We conducted a PRISMA 2020-compliant systematic review and meta-analysis registered with PROSPERO (CRD420251176953). Searches of PubMed, Scopus, Web of Science, and regional repositories (2016–February 2026) identified studies reporting *An. stephensi* bionomics and epidemiological role in Ethiopia. Eligible studies required ≥50% quality score on JBI appraisal tools. Random-effects meta-analysis estimated pooled proportions of *An. stephensi* among total Anopheles, with subgroup analyses by geography, habitat, and behavioural traits. Publication bias was assessed using Egger’s and Begg’s tests.

**Results:** Eighteen studies (9 epidemiological, 11 bionomical) met inclusion criteria. The pooled proportion of *An. stephensi* was 0.51 (95% CI: 0.28–0.75) in epidemiological studies and 0.46 (95% CI: 0.26–0.66) in bionomics studies, with extreme heterogeneity (I² > 99%). Geographic analysis indicated significant variation: south-eastern Ethiopia showed a dominance of 0.73 (95% CI: 0.28–1.18) and eastern Ethiopia 0.57 (95% CI: 0.32–0.82), while central Ethiopia remained lower at 0.13 (95% CI: 0.12–0.14). These findings demonstrate genuine ecological differences rather than methodological objects, with substantial implications for region-specific vector management strategies. Extreme heterogeneity reflected genuine ecological variation across Ethiopia. No evidence of publication bias was detected.

**Conclusion:** *An. stephensi* has very rapidly emerged as a major malaria vector in Ethiopia, with pooled proportions increasing from <10% at first detection (2016) to 51–73% in recent surveys (2024–2025), suggesting ongoing vector displacement parallel to invasion patterns documented in Djibouti. Geographic stratification indicates an urgent need for region-specific urban vector management integrating larval source management, resistance monitoring, and community engagement, particularly in south-eastern Ethiopia, where near-complete vector replacement has occurred.

**Author Summary:** Malaria is usually thought of as a rural disease, but a new mosquito species called *An. stephensi* is changing that picture in Ethiopia. Originally found in South Asia, this mosquito has spread quickly across East Africa and is now common in Ethiopian towns and cities. We reviewed and combined results from published studies to understand how this species behaves and how much it contributes to malaria transmission. Our analysis shows that *An. stephensi* has become one of the dominant malaria vectors in several regions of Ethiopia, especially in the east and south-east, where it has almost replaced other mosquito species. This rapid change means that malaria risk is increasing in urban areas that were previously considered low risk. These findings highlight the urgent need for new control strategies that focus on city environments, such as managing breeding sites, monitoring insecticide resistance, and involving communities in prevention efforts. By understanding how *An. stephensi* is spreading and adapting, we can better protect urban populations and support Ethiopia’s malaria elimination goals.

## Introduction

Malaria remains a significant economic and public health challenge in Ethiopia, which puts millions of people at risk of infection every year [1–4]. Ethiopia’s malaria burden leads to significant economic costs due to healthcare expenses and productivity losses. Traditionally, urban malaria accounted for less than 5% of the national burden, with most cases in rural areas below 2,000 m altitude [2]. However, the emergence of *An. stephensi* poses a risk of concentrating malaria transmission in urban centres, which could intensify due to high population density and overwhelm healthcare systems [3]. This shift may increase the per-capita malaria burden in cities and requires urgent adaptation in control strategies and resource allocation to address the evolving challenge [2–4].

Malaria transmission in the country has historically been dominated by the vector *An. arabiensis*, with secondary vectors such as *Anopheles funestus, Anopheles pharoensis*, and *Anopheles nili,* as well as *Anopheles coustani* [5–6]. The recent invasion of *An. stephensi*, a vector previously confined to South Asia and the Middle East, has complicated the Ethiopian malaria epidemiology [7, 8]. Unlike traditional rural vectors, *An. stephensi* exhibits ecological adaptation, thriving in urban settings using artificial water sources, including overhead tanks, construction sites, and waste containers [9, 10]. This adaptability allows it to colonise densely populated areas, challenging the long-held assumption that urbanisation reduces malaria risk [11, 12]. Its capacity to transmit both *Plasmodium falciparum* and *Plasmodium vivax* complicates the challenge, as Ethiopia is currently dealing with a dual burden of these two parasites. This indicates that the first invasive malaria vector has established itself in urban Africa, suggesting a paradigm shift in transmission dynamics [8, 12].

*An. stephensi’* historical distribution was mostly limited to South Asia, particularly India, Pakistan, and parts of the Arabian Peninsula [8–14]. Its detection in Djibouti in 2012 marked the beginning of its rapid expansion across East Africa [15, 16]. The invasion of *An. stephensi* across East Africa follows a predictable ecological expansion pattern. Initially detected in Djibouti in 2012, the vector rapidly achieved 80–95% dominance within 5–8 years, displacing native Anopheles species and prompting malaria outbreak responses [7, 10, 11, 15, 16]. In Ethiopia, its first detection in Dire Dawa in 2016 signalled the start of its documented expansion. By 2017–2018, the species was verified in the Somali and Afar regions, with reports from 2019–2025 indicating ongoing geographic expansion, resulting in near-total dominance (88–99% of Anopheles populations) in south-eastern Ethiopia [15–16]. This invasion trajectory indicates that Ethiopia could experience complete native vector displacement by 2028–2032, significantly altering the country’s malaria epidemiology within the next 5–10 years [19–20].

Subsequent reports confirmed its presence in Ethiopia in 2016, first in Dire Dawa and later expanding to several regions, including Somali, Afar, and, most recently southern Ethiopia [17, 18, 21, 25]. Its expansion correlates with growing urbanisation, which produces favourable breeding conditions. Alarmingly, modelling studies suggest that the spread of *An. stephensi* might put over 126 million Africans at risk for malaria [19]. Its invasion represents not only geographical expansion but also a shift in malaria transmission, particularly in previously low-prevalence urban areas. *An. stephensi* role in urban malaria is becoming increasingly evident, with its prevalence associated with increased malaria incidence in Ethiopian urban areas. Unlike rural vectors, *An. stephensi* bites early in the evening, reducing the effectiveness of long-lasting insecticidal nets (LLINs) [21, 27]. This behavioural shift requires a re-evaluation of current control strategies, including LLINs and IRS [27].

The adaptability across ecological niches, as well as the increasing resistance to widely used insecticides, highlights its importance in changing the epidemiology of malaria in Ethiopia [22, 23]. Because the species preferentially uses human-made habitats, traditional vector management strategies based on rural transmission may not be sufficient to address the challenges posed by this invasive vector. Failure to adapt policies could reverse two decades of malaria control progress [24]. The emerging evidence shows that *An. stephensi* may influence the seasonal malaria transmission patterns, potentially extending transmission periods in urban centres [23].

Ethiopia has made significant achievements in decreasing malaria incidence during the last two decades, but *the An. stephensi* invasion threatens to undermine these successes [25]. The World Health Organization (WHO) has identified *An. stephensi* as a significant new vector in Africa and warns that its spread might threaten malaria control efforts across the continent [13]. Ethiopia’s growing urbanisation, which provides ideal habitats, makes resolving this issue even more urgent. This challenge is not confined to Ethiopia; neighbouring East African countries face similar risks, showing the continental implications of *An. stephensi* spread [13, 25].

Despite increasing indications of *An. stephensi* spreading, significant gaps remain in the understanding of its biology, epidemiology, and resistance patterns in Ethiopia. Studies have documented its spatiotemporal distribution and seasonal dynamics, as well as its establishment in various ecological zones [22, 25], but a comprehensive review is required. Resistance to pyrethroids and carbamates affects the efficacy of LLINs and indoor residual spraying (IRS) [26, 27]. Without a complete synthesis, control programmes risk depending on fragmented evidence. This review therefore presents the first pooled quantitative synthesis of *An. stephensi* bionomics and epidemiology in Ethiopia.

Despite increased studies on *An. stephensi* in Africa, significant evidentiary gaps remain, particularly regarding its emergence in Ethiopia. Although *An. stephensi* has been identified as a novel vector danger and the Horn of Africa is receiving increased attention, there is currently no complete quantitative synthesis to guide Ethiopia’s National Malaria Elimination Program. Regional systematic reviews include larger East African epidemiology, but they lack the geographic differentiation and thorough bionomic characterisation required for evidence-based urban vector management at the regional and district levels. This review covers that crucial vacuum by integrating all available quantitative data on *An. stephensi* distribution, bionomics, resistance patterns, and epidemiological importance across Ethiopia’s varied ecological zones.

Historically, Ethiopia’s urban malaria control strategies have focused on reducing habitats and promoting the use of LLINs as a control method. However, *An. stephensi* ecology, including its preference for artificial containers and biting behaviour, makes these measures ineffective [25, 28, 20]. Insecticide resistance complicates control, demanding new approaches including larval source management, integrated vector control, and community-based interventions [26]. Findings will directly inform Ethiopia’s National Malaria Elimination Program and WHO regional vector control frameworks.

This systematic review addresses this gap by synthesising all available quantitative data to provide policymakers and control programmes with an evidence-based understanding of *An. stephensi* establishment and invasive potential in Ethiopia. It aims to determine the distribution and proportion of *An. stephensi* within Ethiopian Anopheles populations, evaluate its epidemiological importance across various geographic and ecological settings, and characterise the bionomic traits. The seasonal variations, evidence quality, and research gaps were examined with contrasts to other East African countries where *An. stephensi* has been identified. By combining various findings, this review gives a foundation for designing context-specific, evidence-based interventions against *An. stephensi* in Ethiopia and beyond.

## Method

### Design and Protocol

This systematic review and meta-analysis followed the Preferred Reporting Items for Systematic Reviews and Meta-Analyses Protocol (PRISMA-P 2020) guidelines [29]. The study selection process was documented with a four-stage PRISMA flow diagram, which depicts the steps taken from the initial collection of identified records to the final set of studies included in the analysis. The review protocol was developed in advance to ensure methodological transparency, and it was formally registered with the International Prospective Register of Systematic Reviews (PROSPERO) under the registration number CRD420251176953.

### Database and Search Strategy

This systematic review and meta-analysis, which started on November 30, 2025, focused on articles from Ethiopia published in English until February 10, 2026. A detailed analysis of electronic databases and grey literature was carried out, including searches in PubMed, Scopus, Science Direct, Web of Science, Google Scholar, and regional repositories. The selected databases were chosen for their relevant literature indexing, and additional local research was collected via Google Scholar, despite its systematic review constraints. Before screening the title and abstract, duplicate records were excluded with EndNote X7.

Search terms were combined with Boolean operators (“AND” and “OR”) and adapted for each platform’s syntax. The primary search string used across databases was (’Anopheles stephensi’ OR ‘urban mosquito’) AND (Ethiopia OR ‘eastern’ OR ‘southern’ OR ‘central’ OR ‘south-western’) AND (’bionomics’ OR ‘epidemiology’ OR ‘host preference’ OR ‘resting behaviour’ OR ‘breeding habitat’ OR ‘distribution’ OR ‘malaria’). The search for *An. stephensi* in Dire Dawa, Ethiopia, began in January 2016, coinciding with its first finding. There is no preceding research, allowing us to focus on post-invasion data while ignoring pre-invasion literature. This search will finish on February 10, 2026, using parameters for publication date (2016-February 2026) and language (English). Additional relevant studies were discovered by reviewing reference lists using a snowballing method.

### Eligibility Criteria

Records were imported into EndNote X7 for organization and duplicate removal. Inclusion criteria were: studies conducted in Ethiopia; entomological data on Anopheles (any life stage); outcomes reporting the proportion of *An. stephensi* or bionomic traits (habitat, feeding, resting, behaviour); and study designs including cross-sectional, case-control, cohort/prospective, modelling studies, and systematic reviews with primary data. Exclusion criteria were: studies outside Ethiopia; human-only clinical studies without entomological data; case reports, editorials, opinion pieces, and conference abstracts without full text; non-English publications; and studies published before 2016. Studies were eligible only if they scored ≥50% on the JBI quality appraisal. The eligibility criteria for the studies were shown in table 1.

**Table 1.**
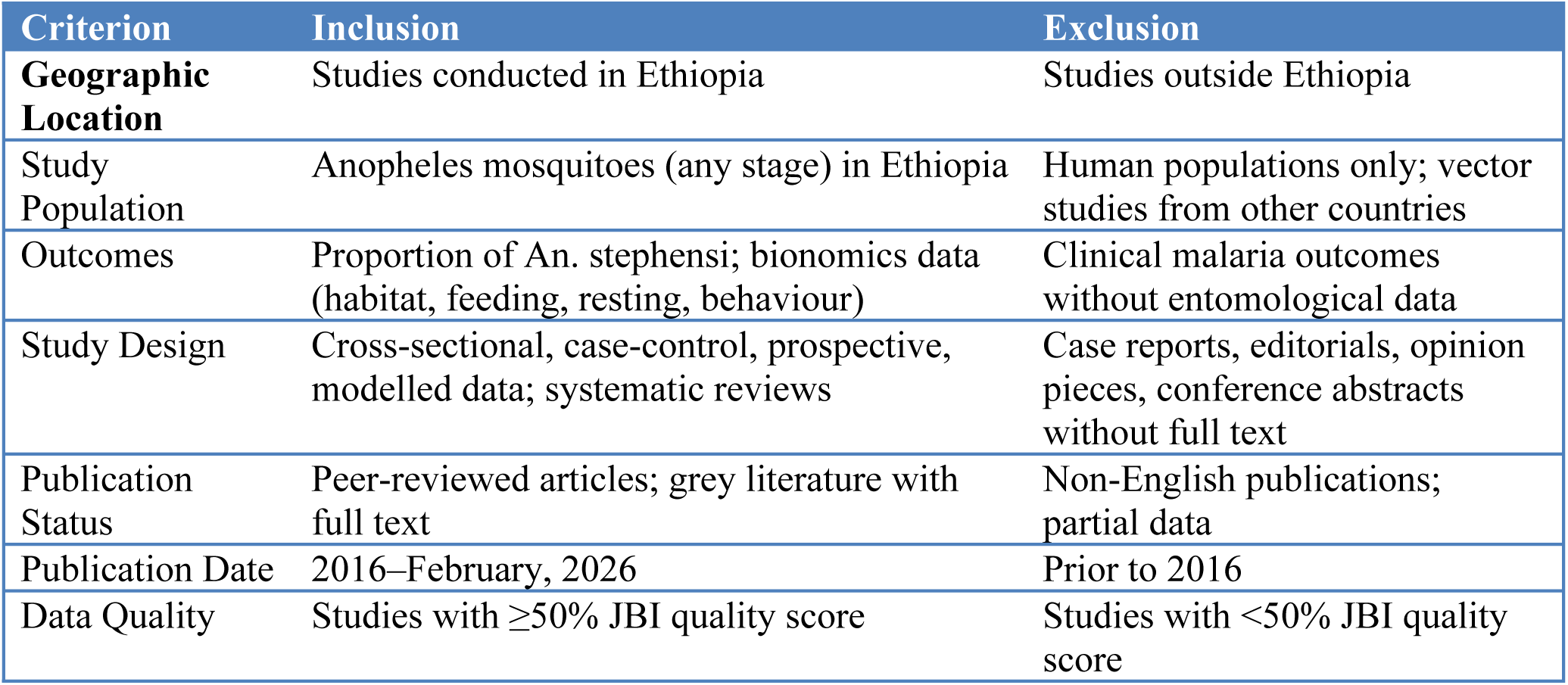
summary inclusion and exclusion criteria.

| Criterion | Inclusion | Exclusion |
| --- | --- | --- |
| **Geographic Location** | Studies conducted in Ethiopia | Studies outside Ethiopia |
| Study Population | Anopheles mosquitoes (any stage) in Ethiopia | Human populations only; vector studies from other countries |
| Outcomes | Proportion of *An. stephensi*; bionomics data (habitat, feeding, resting, behaviour) | Clinical malaria outcomes without entomological data |
| Study Design | Cross-sectional, case-control, prospective, modelled data; systematic reviews | Case reports, editorials, opinion pieces, conference abstracts without full text |
| Publication Status | Peer-reviewed articles; grey literature with full text | Non-English publications; partial data |
| Publication Date | 2016–February, 2026 | Prior to 2016 |
| Data Quality | Studies with $\geq 50\%$ JBI quality score | Studies with $< 50\%$ JBI quality score |

### Outcome of Interest

The primary outcome was the proportion of *An. stephensi* among all Anopheles collected in each study, expressed as n/N. Secondary outcomes included bionomic characteristics (breeding habitat type, host preference, resting location, and feeding time), seasonal patterns, surveillance method, and ecological zone.

### Study Selection and quality Assessment

Two reviewers (T.B. and K.L.) independently reviewed titles and abstracts. Then, three reviewers (T.B., K.L., and A.D.) assessed full-text publications for eligibility criteria. Disagreements were resolved by discussion and, where necessary, adjudicated by a fourth reviewer (L.R.). Study quality was assessed using the Joanna Briggs Institute (JBI) Critical Appraisal Checklist for Prevalence Studies for cross-sectional designs and the JBI Checklist for Cohort Studies for prospective designs. Each item was scored as “yes” (1), “no” (0), or “unclear” (0), yielding a maximum score of 9. Studies scoring ≥4.5 (50%) were included (30).

### Data Extraction

A standardised data extraction form was tested on three studies and then applied to all included studies. Extracted items included first author, year, study location and ecological zone, study design, sampling method, sample frame, collection period and season, total Anopheles collected (N), number of *An. stephensi* (n), method of species identification (morphological and/or molecular), and bionomic variables (breeding habitat, host preference, resting behaviour, and feeding time). Funding sources and declared conflicts of interest were also recorded. Two reviewers independently extracted data; discrepancies were resolved by consensus.

### Data Synthesis and Analysis

All extracted data was organised in Excel and then imported into STATA version 19.5 (StataCorp, College Station, Texas, USA) for statistical analysis. A random-effects meta-analysis approach was used to obtain pooled estimates of *An. stephensi* proportions, taking into account study heterogeneity. The subgroup analyses were established to investigate potential sources of heterogeneity based on ecological factors (geographic zone, habitat type), vector behaviour (host preference, resting patterns, feeding time), and study design factors (epidemiological domain, surveillance method, collection season). Significant subgroup differences were found using interaction testing (Q statistic) at p < 0.05. The point estimates were reported with 95% confidence intervals. The I² statistic was used to quantify heterogeneity, with the following interpretations: I² < 25% = low, 25-50% = moderate, 50-75% = large, and > 75% = extreme heterogeneity. Random-effects models were used throughout, with the expectation of significant heterogeneity due to regional and ecological variation. Tau² estimations are presented alongside I² to measure the variance between studies. Publication bias was evaluated using (1) visual assessment of the funnel plot; (2) Egger’s weighted regression test (p < 0.05 showing bias); (3) Begg’s rank correlation test (p < 0.05 indicating bias); and (4) trim-and-fill analysis to estimate the number and size of potentially missing studies. Stata’s metabias and metatrim commands were used.

## Results

### Search Results

A total of 4,410 records were identified from electronic databases, including PubMed (2,892 records), Science Direct (365 records), and Scopus (218 records), as well as an additional 752 records from Google Scholar. After eliminating 2,832 duplicate records, 2,393 records remained for screening. During the screening stage, 2,017 records were excluded after a review of their titles and abstracts. A full-text screening of 107 publications showed 49 articles that were potentially relevant based on title and abstract review. During the full-text review, 33 papers were removed for the following reasons: lack of outcome of interest (n = 24), the study location being outside Ethiopia (n = 4), non-English publishing (n = 3), and insufficient methodological transparency (n = 2). This identified 18 eligible studies for inclusion in the systematic review. As shown in Figure 1, the PRISMA 2020 flow diagram summarizes the study selection process and reasons for exclusion.

**Figure 1.**
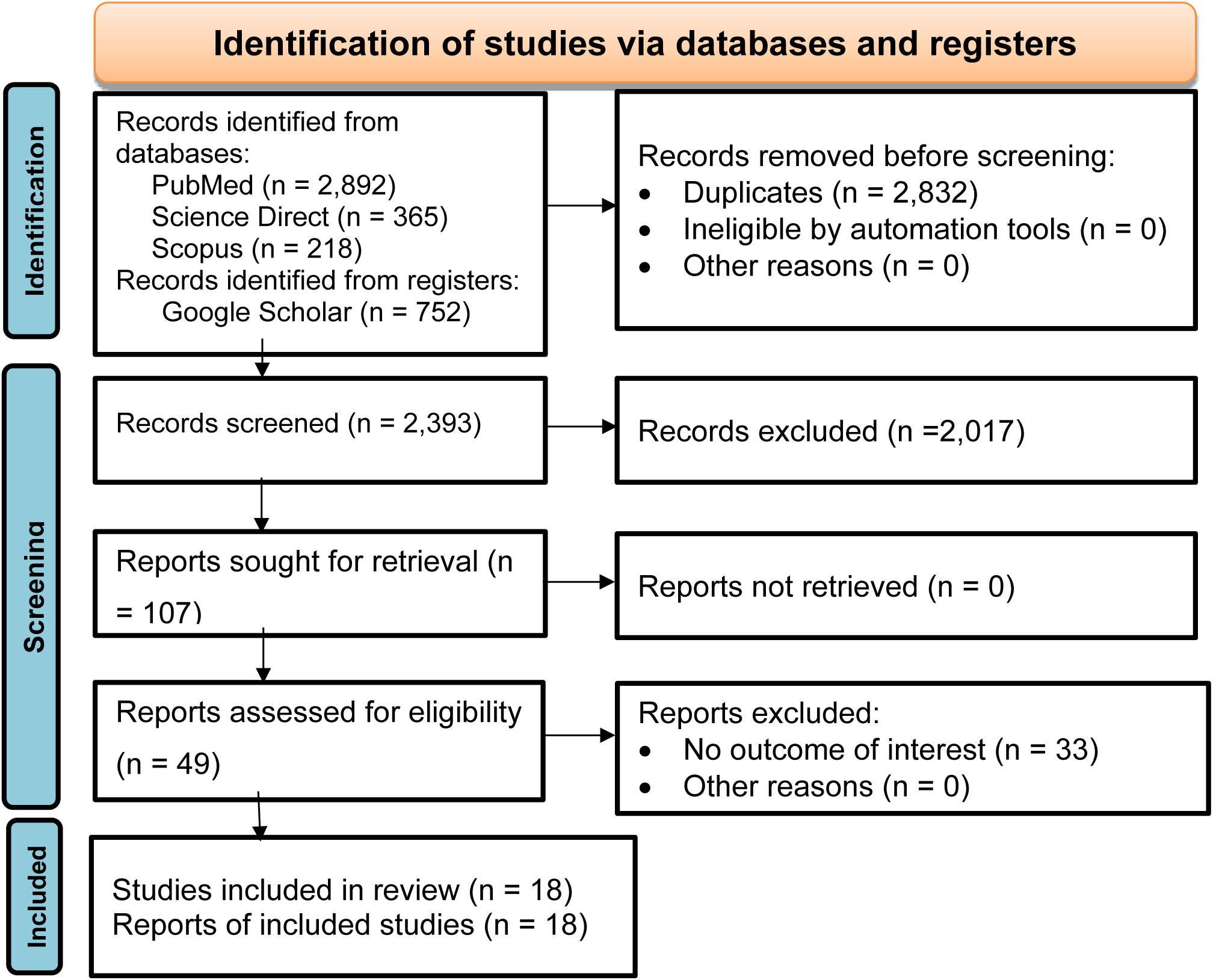
PRISMA 2020 flow diagram showing the identification, screening, eligibility assessment, and inclusion of studies in the systematic review

### Characteristics of the Included Studies

A total of 18 studies published between 2016 and 2025 were analysed [20, 25, 28, 31, 32, 33, 34, 35, 36, 37, 38, 39, 40, 41, 42, 43, 44, 45]. Most of the studies were conducted in Eastern Ethiopia (n=12), with additional studies from Central Ethiopia (n=2), Southern Ethiopia (n=1), and South-eastern Ethiopia (n=2), along with one multi-site study. Sample sizes varied significantly, from a low of 34 specimens [31] to a high of 49,482 specimens [32], with reported proportions of *An. stephensi* values ranging from 0.07 to 1.0. Several studies [33–36] reported very high proportions (≥0.8), particularly in Eastern Ethiopia. The studies reviewed showed considerable variability in sample sizes, ranging from 34 to 49,482 participants. A significant 67% of the studies were published between 2023 and 2025, suggesting increased research output following the emergence of *An. stephensi*. Moreover, geographic representation revealed that 67% of the studies were focused in Eastern Ethiopia, the original detection site of *An. stephensi*, indicating notable research gaps in Central and South-western Ethiopia. Quality appraisal scores ranged from 7 to 9, with most of the studies scoring 8 or 9, strengthening confidence in pooled estimates. A detailed summary of included studies was pshown in the below Table 2.

**Table 2.**
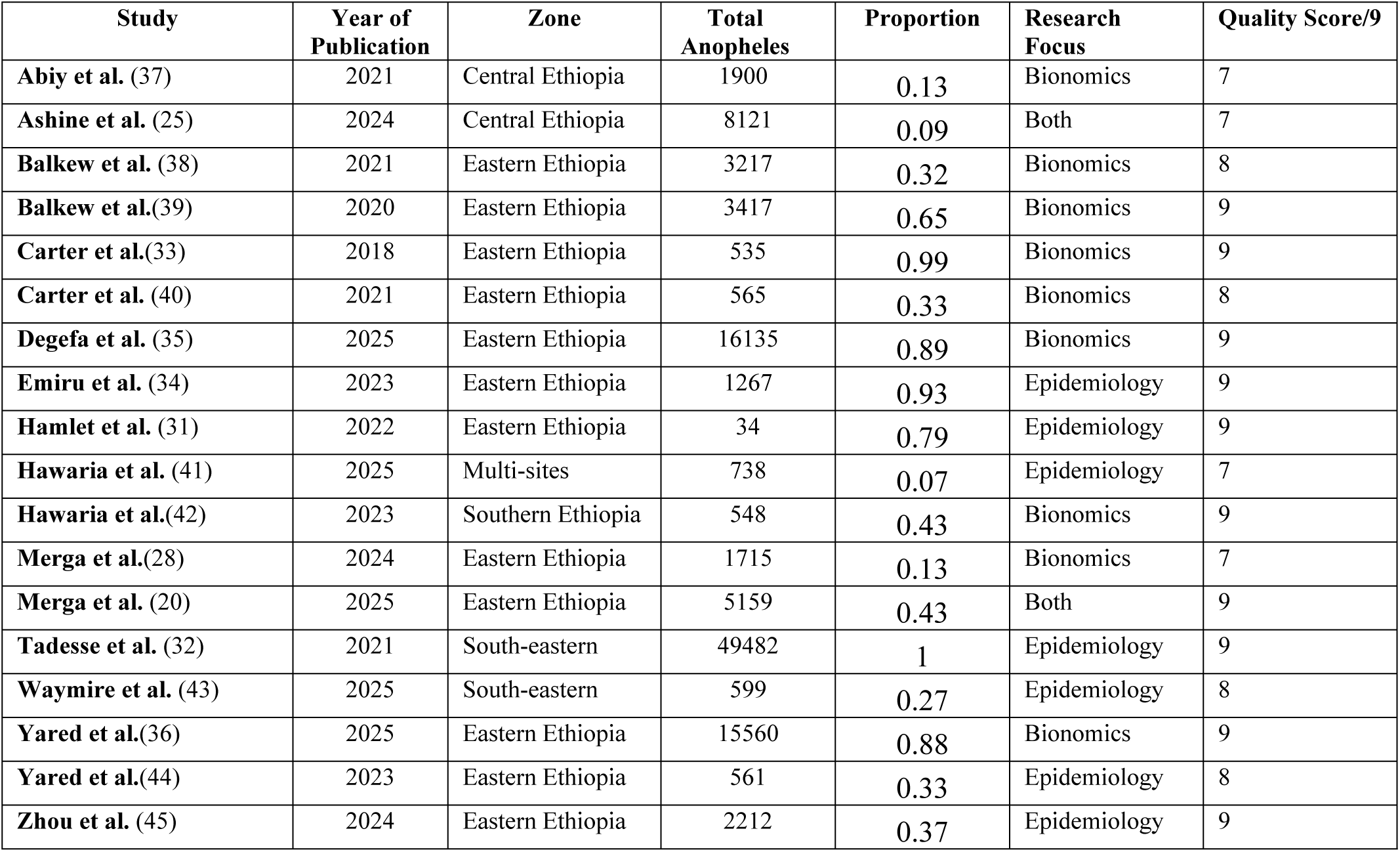
Characteristics of included studies on *An. stephensi* in Ethiopia 2016–2025 Both = studies reporting both epidemiological and bionomic outcomes.

| Study | Year of Publication | Zone | Total Anopheles | Proportion | Research Focus | Quality Score/9 |
| --- | --- | --- | --- | --- | --- | --- |
| Abiy et al. (37) | 2021 | Central Ethiopia | 1900 | 0.13 | Bionomics | 7 |
| Ashine et al. (25) | 2024 | Central Ethiopia | 8121 | 0.09 | Both | 7 |
| Balkew et al. (38) | 2021 | Eastern Ethiopia | 3217 | 0.32 | Bionomics | 8 |
| Balkew et al.(39) | 2020 | Eastern Ethiopia | 3417 | 0.65 | Bionomics | 9 |
| Carter et al.(33) | 2018 | Eastern Ethiopia | 535 | 0.99 | Bionomics | 9 |
| Carter et al. (40) | 2021 | Eastern Ethiopia | 565 | 0.33 | Bionomics | 8 |
| Degefa et al. (35) | 2025 | Eastern Ethiopia | 16135 | 0.89 | Bionomics | 9 |
| Emiru et al. (34) | 2023 | Eastern Ethiopia | 1267 | 0.93 | Epidemiology | 9 |
| Hamlet et al. (31) | 2022 | Eastern Ethiopia | 34 | 0.79 | Epidemiology | 9 |
| Hawaria et al. (41) | 2025 | Multi-sites | 738 | 0.07 | Epidemiology | 7 |
| Hawaria et al.(42) | 2023 | Southern Ethiopia | 548 | 0.43 | Bionomics | 9 |
| Merga et al.(28) | 2024 | Eastern Ethiopia | 1715 | 0.13 | Bionomics | 7 |
| Merga et al. (20) | 2025 | Eastern Ethiopia | 5159 | 0.43 | Both | 9 |
| Tadesse et al. (32) | 2021 | South-eastern | 49482 | 1 | Epidemiology | 9 |
| Waymire et al. (43) | 2025 | South-eastern | 599 | 0.27 | Epidemiology | 8 |
| Yared et al.(36) | 2025 | Eastern Ethiopia | 15560 | 0.88 | Bionomics | 9 |
| Yared et al.(44) | 2023 | Eastern Ethiopia | 561 | 0.33 | Epidemiology | 8 |
| Zhou et al. (45) | 2024 | Eastern Ethiopia | 2212 | 0.37 | Epidemiology | 9 |

### Epidemiological proportion of *Anopheles stephensi*

The meta-analysis included nine studies focusing on the epidemiological proportion of *An. stephensi* among Anopheles mosquitoes in diverse ecological and urban contexts in Ethiopia. The reported proportions varied widely, from 0.07 in [41] and 0.08 in [46] to near-total dominance of 1.00 [32]. Very high proportions of *An. stephensi* have been documented with proportions of 0.93 [34], 0.79 [31], and 0.79[47]. Moderate proportions of ecological variables were observed [45] with a reported estimate of 0.37 [44] at 0.33 and [43] at 0.27, indicative of mixed-species habitats or transitional ecological zones. The overall pooled proportion was 0.51 (95% CI: 0.28-0.75; z = 4.24, *p* < 0.001). Heterogeneity was extreme (τ² = 0.13, I² = 99.98%, H² = 4393.96), reflecting genuine ecological and epidemiological variability across Ethiopia. This underscores the importance of stratified analyses by ecological zone. Results are summarised in Figure 2.

**Figure 2.**
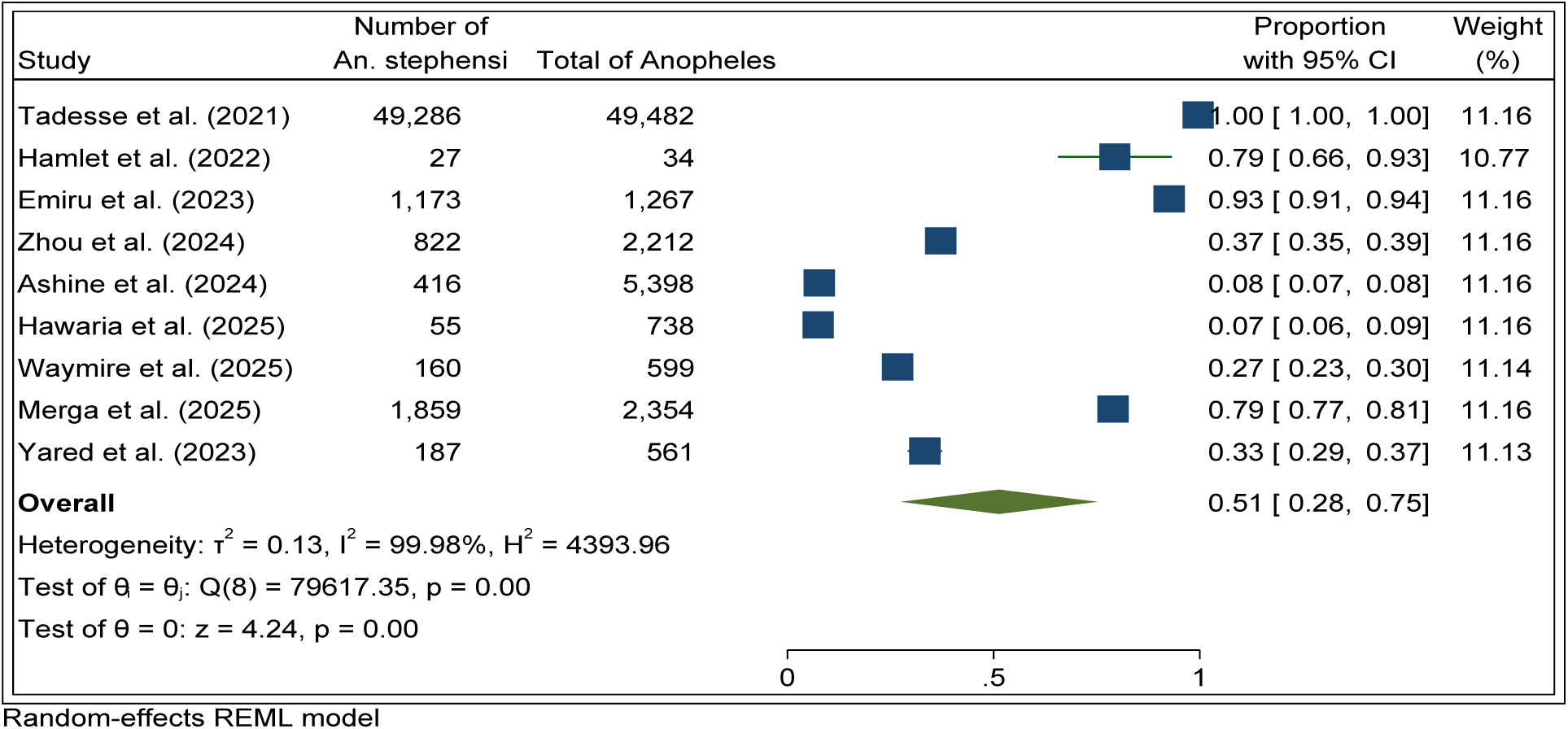
Forest plot of pooled epidemiological proportion of *An. stephensi* among total *Anopheles* mosquitoes in Ethiopia.

### Subgroup Analysis by Geographic Zone and Epidemiological Domain

Subgroup analysis by geographic zone showed significant variation in the proportion of *An. stephensi* among all *Anopheles* mosquitoes across Ethiopia. In Eastern Ethiopia, the pooled proportion was 0.57 (95% CI: 0.32–0.82), with extreme heterogeneity (τ² = 0.06, I² = 99.69%). Estimates ranged from 0.33 [44] to 0.79 [31, 20]. In multi-site studies, the pooled proportion was much lower at 0.08 (95% CI: 0.07–0.08), with negligible heterogeneity (I² = 0.07%), suggesting consistent results across ecological zones [25, 41]. The geographic heterogeneity in *An. stephensi* proportions across Ethiopia (Eastern 0.57, Central 0.13, and South-eastern 0.73) indicates significant ecological and epidemiological differences as shown by a significant Q-statistic (Q(2) = 22.89, p < 0.001).

Central Ethiopia’s low proportion relates to its cooler highland climate (1,500–2,500 metres), which may limit larval development. In contrast, Eastern Ethiopia’s moderate-high dominance corresponds to semi-arid lowlands with consistent breeding conditions, while South-eastern Ethiopia’s high dominance is linked to urbanization and favourable breeding environments created by infrastructure development. In South-eastern Ethiopia, the pooled proportion reached 0.73 (95% CI: 0.28–1.18), with marked heterogeneity (τ² = 0.16, I² = 99.97%), largely driven by differences between [32] and [43]. Overall, the pooled proportion across all zones was 0.51 (95% CI: 0.28–0.75), and subgroup differences were statistically significant (Q(2) = 22.89, p < 0.001; Figure 3).

**Figure 3.**
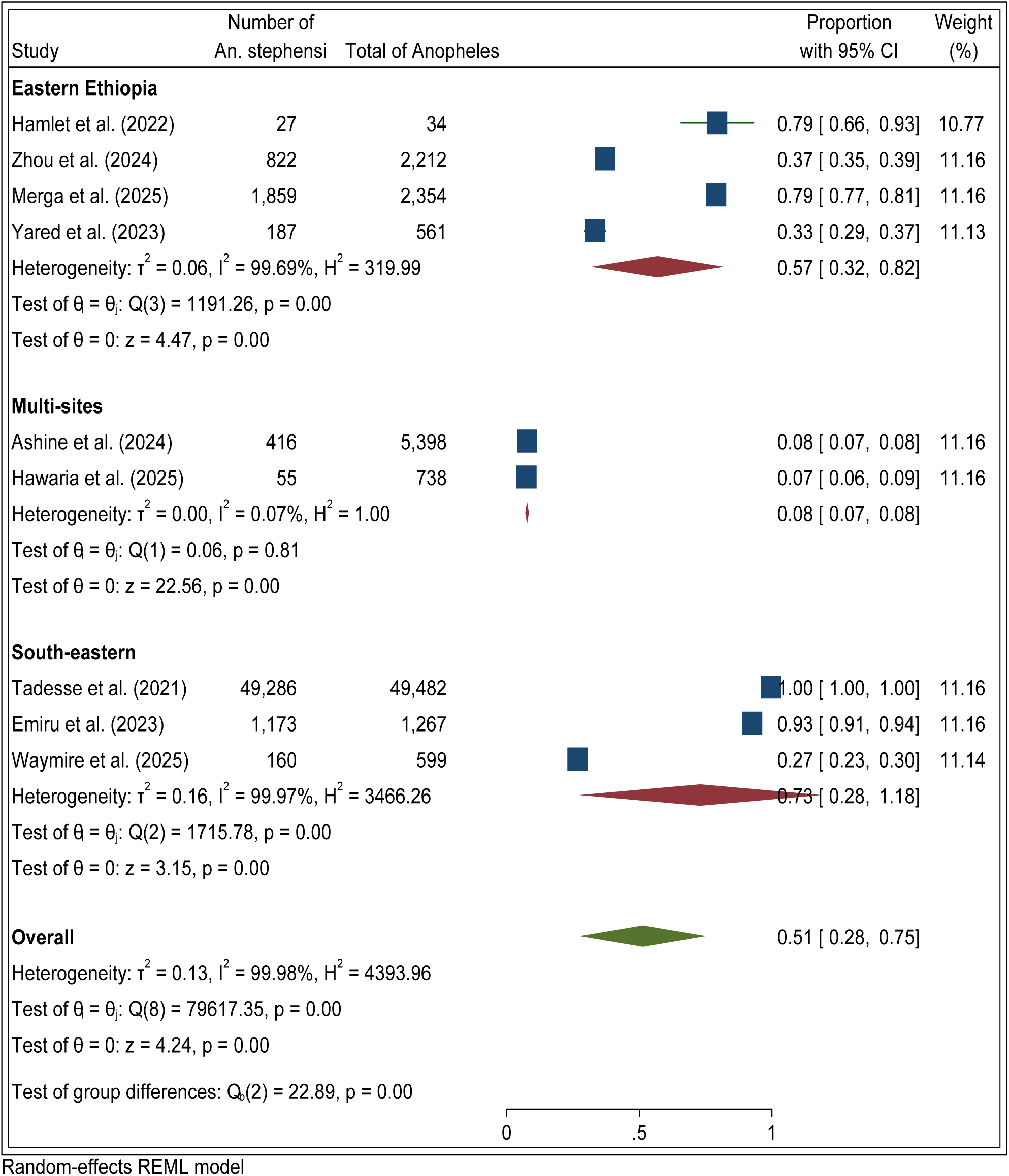
Forest plot of subgroup meta-analysis of *An. stephensi* proportions by geographic zone in Ethiopia

Similarly, subgroup meta-analysis across epidemiological domains showed distinct trends. In the entomological + molecular category, proportions ranged from 0.07 [41] to 1.00 [32], with a pooled estimate of 0.35 (95% CI: 0.08–0.78; z = 1.62, p = 0.11) and high heterogeneity (τ² = 0.19, I² = 99.99%). The epidemiology group included two studies, with proportions ranging from 0.37 [45] to 0.93 [46]; the pooled estimate was 0.65 (95% CI: 0.11–1.19; z = 2.34, p = 0.02), with marked heterogeneity (τ² = 0.15, I² = 99.95%). The modelling group reported proportions from 0.33 [44] to 0.79 [31, 20], yielding a pooled estimate of 0.64 (95% CI: 0.33–0.94; z = 4.12, p < 0.001) with high heterogeneity (τ² = 0.07, I² = 99.39%). The overall pooled proportion across all domains was 0.51 (95% CI: 0.28–0.75), and subgroup differences were not statistically significant (Q(2) = 1.23, p = 0.54; Figure 4).

**Figure 4.**
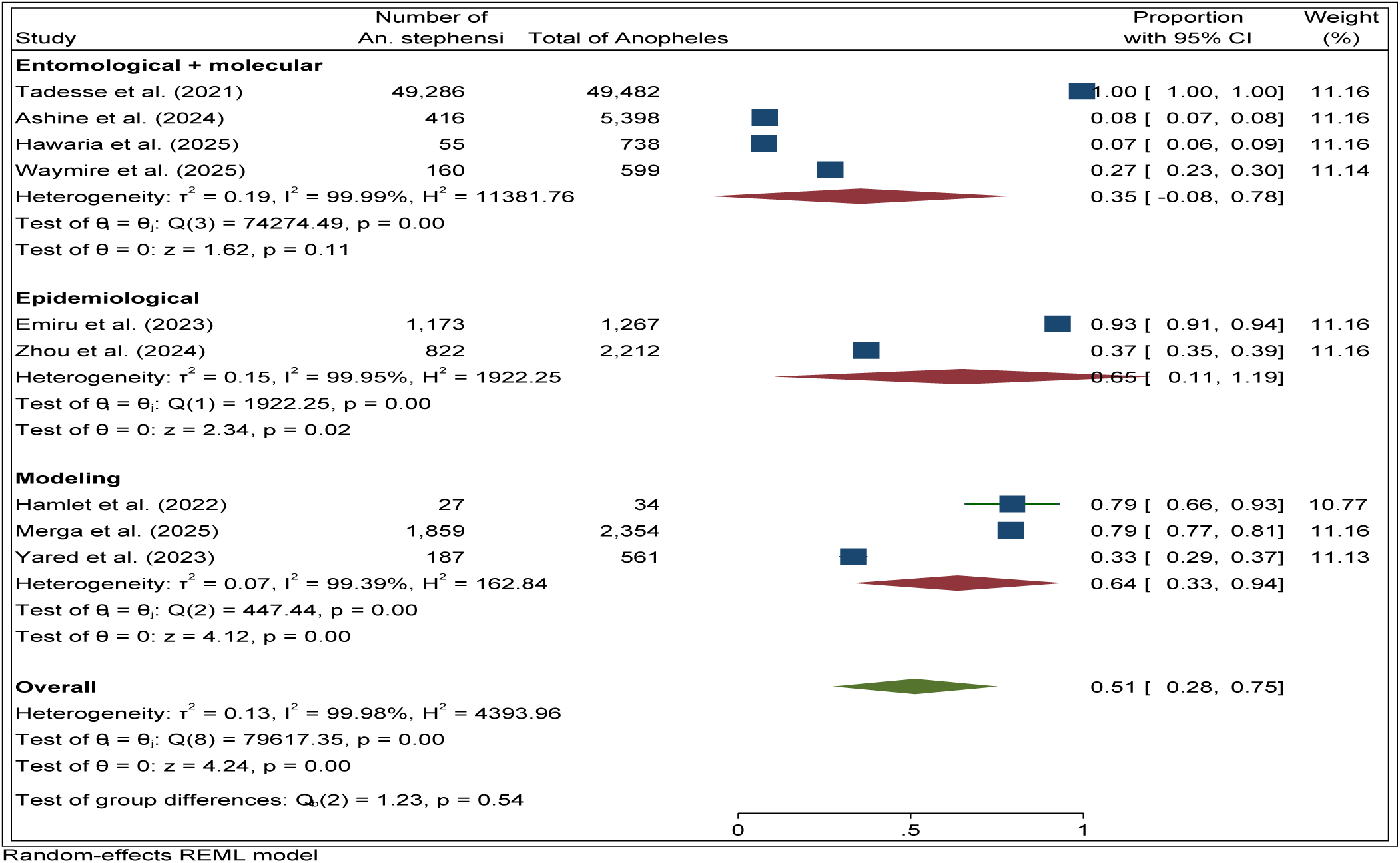
Forest plot of subgroup meta-analysis of *An. stephensi* proportions by epidemiological domain (entomological + molecular, epidemiology, modelling)

### Subgroup analysis by study design

Subgroup meta-analysis across epidemiological study designs showed distinct trends in the proportion of *An. stephensi* among total *Anopheles* mosquitoes. The cross-sectional studies reported proportions ranging from 0.07 [41] to 1.00 [32], with a pooled estimate of 0.35 (95% CI: 0.08–0.78; z = 1.62, p = 0.11). Heterogeneity was extreme (τ² = 0.19, I² = 99.99%). The modelling group included studies [8, 20, 44] with proportions ranging from 0.33 to 0.79, with a pooled estimate of 0.64 (95% CI: 0.33–0.94; z = 4.12, p < 0.001) and high heterogeneity (τ² = 0.07, I² = 99.39%). Prospective case–control studies [34, 45] reported proportions of 0.93 and 0.37 and a pooled estimate of 0.65 (95% CI: 0.11–1.19; z = 2.34, p = 0.02) and marked heterogeneity (τ² = 0.15, I² = 99.95%). The overall pooled proportion across all study designs was 0.51 (95% CI: 0.28–0.75), with extreme heterogeneity (I² = 99.98%). Subgroup differences were not statistically significant (Q(2) = 1.23, p = 0.54; Figure 5).

**Figure 5.**
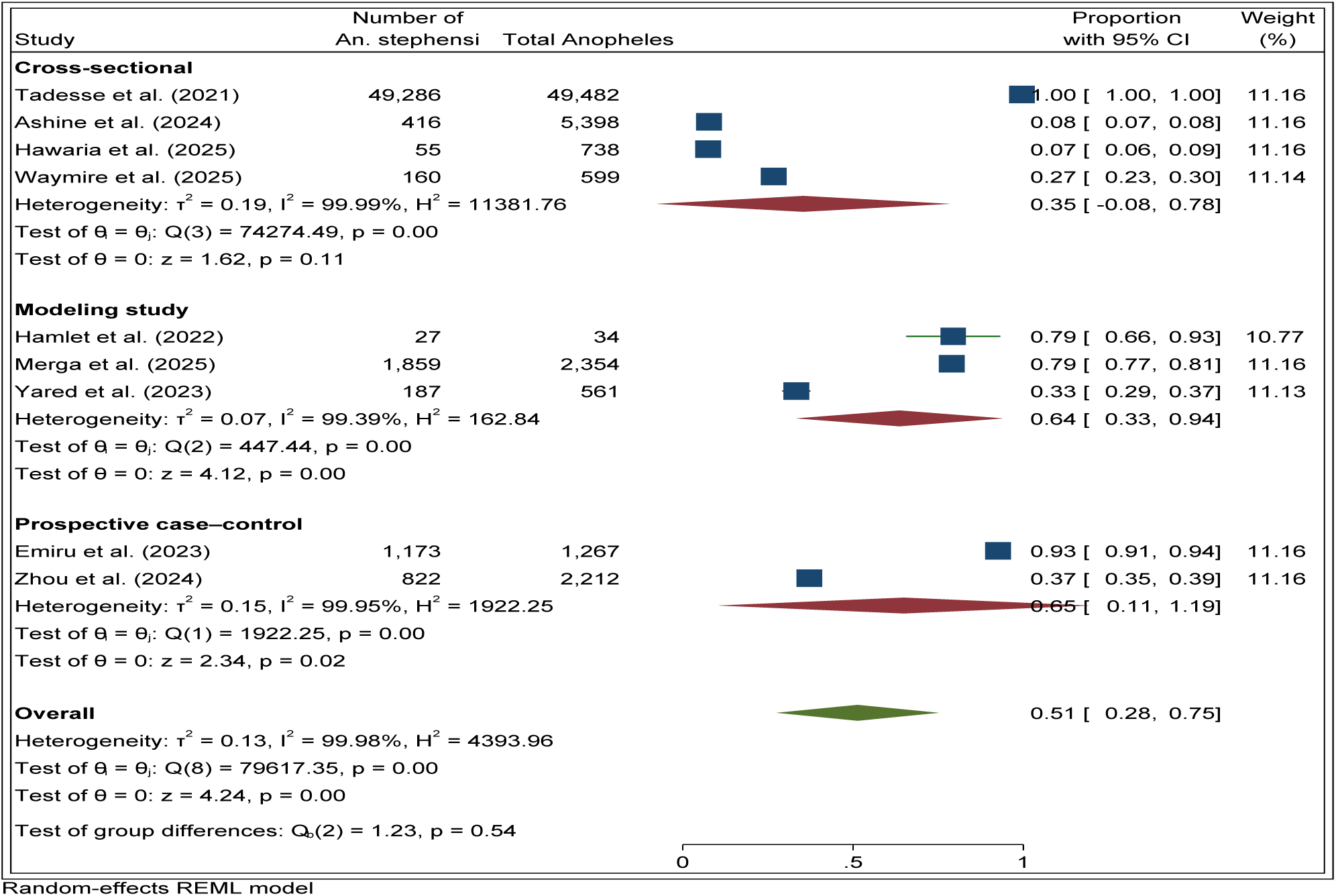
Forest plot of subgroup meta-analysis of *An. stephensi* proportions by study design (cross sectional, modelling, case control)

### Funnel Plot and Publication Bias Assessment

The funnel plot analysis showed no evidence of publication bias in the epidemiological dataset. The inverse-variance weighted estimate (θ_IV) indicated central alignment and symmetrical distribution of studies within the pseudo 95% confidence limits (Figure 6). Egger’s regression test (β₁ = 2.67, p = 0.685) and Begg’s test (Kendall’s score = 2.00, p = 0.917) were non-significant. Trim-and-fill analysis confirmed symmetry, with no imputed studies and a stable pooled estimate (0.989, 95% CI: 0.988–0.989). This suggests robustness of the pooled results despite high heterogeneity.

**Figure 6.**
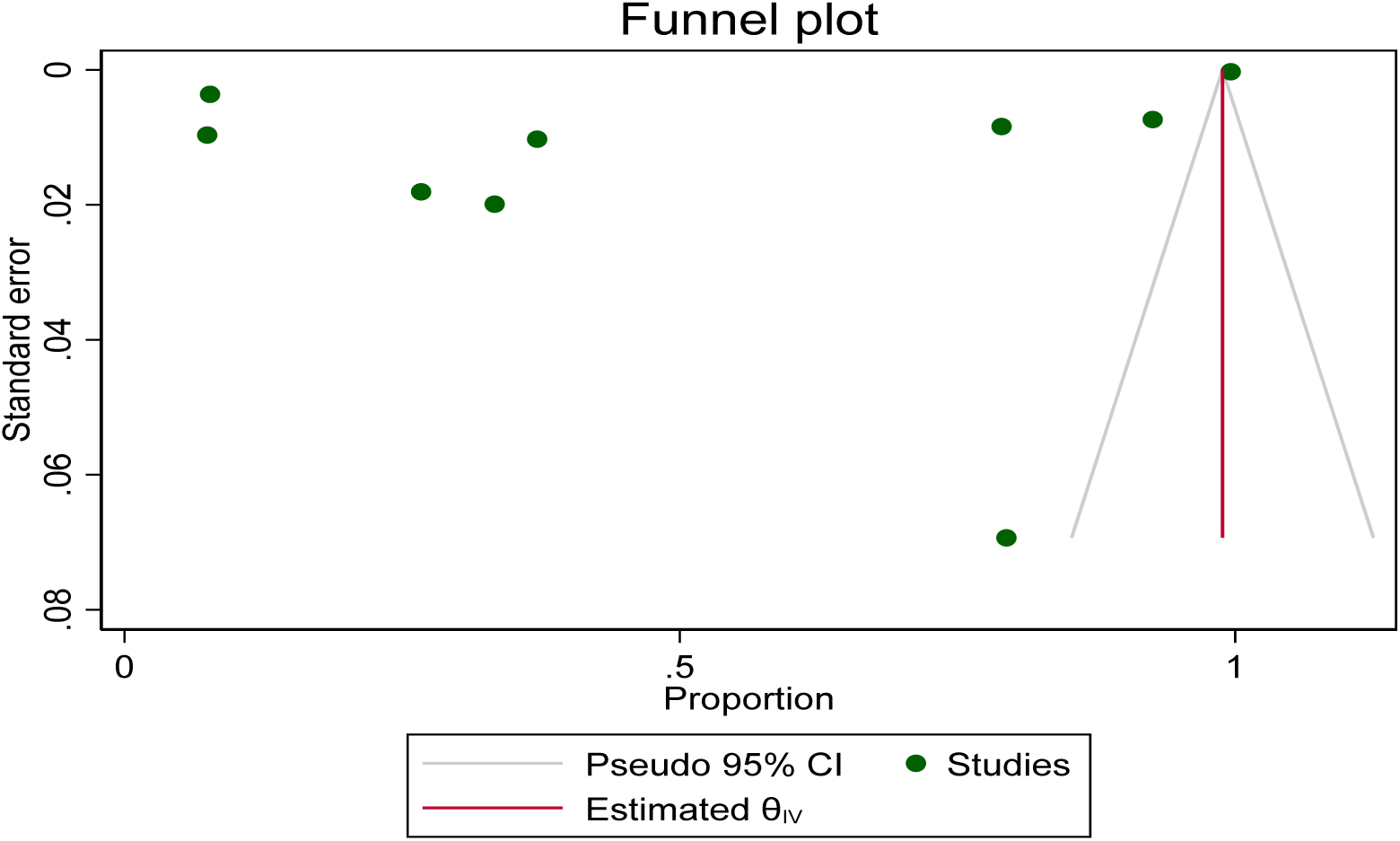
Funnel plot assessing publication bias in epidemiological studies of *An. stephensi* in Ethiopia

### Proportion of *Anopheles stephensi* Bionomics

The forest plot meta-analysis of bionomics studies revealed substantial variation in the proportion of *An. stephensi* across ecological and behavioural contexts. Estimates ranged from a low of 0.13 [46, 28, 37] to a high of 0.99 [48]. Intermediate proportions were reported at 0.32 [38], 0.33 [40], and 0.43 [18]. Degefa et al. [35] reported a proportion of 0.89, closely followed by Yared et al. [36] with 0.88, both supported by large sample sizes. The random-effects model yielded a pooled proportion of 0.46 (95% CI: 0.26–0.66; z = 4.46, p < 0.001). Heterogeneity was extreme (τ² = 0.11, I² = 99.98%, H² = 4,120.87), reflecting ecological diversity and methodological variability across studies. This confirms the significant presence of *An. stephensi* in diverse bionomic contexts. Results are summarised in Figure 7.

**Figure 7.**
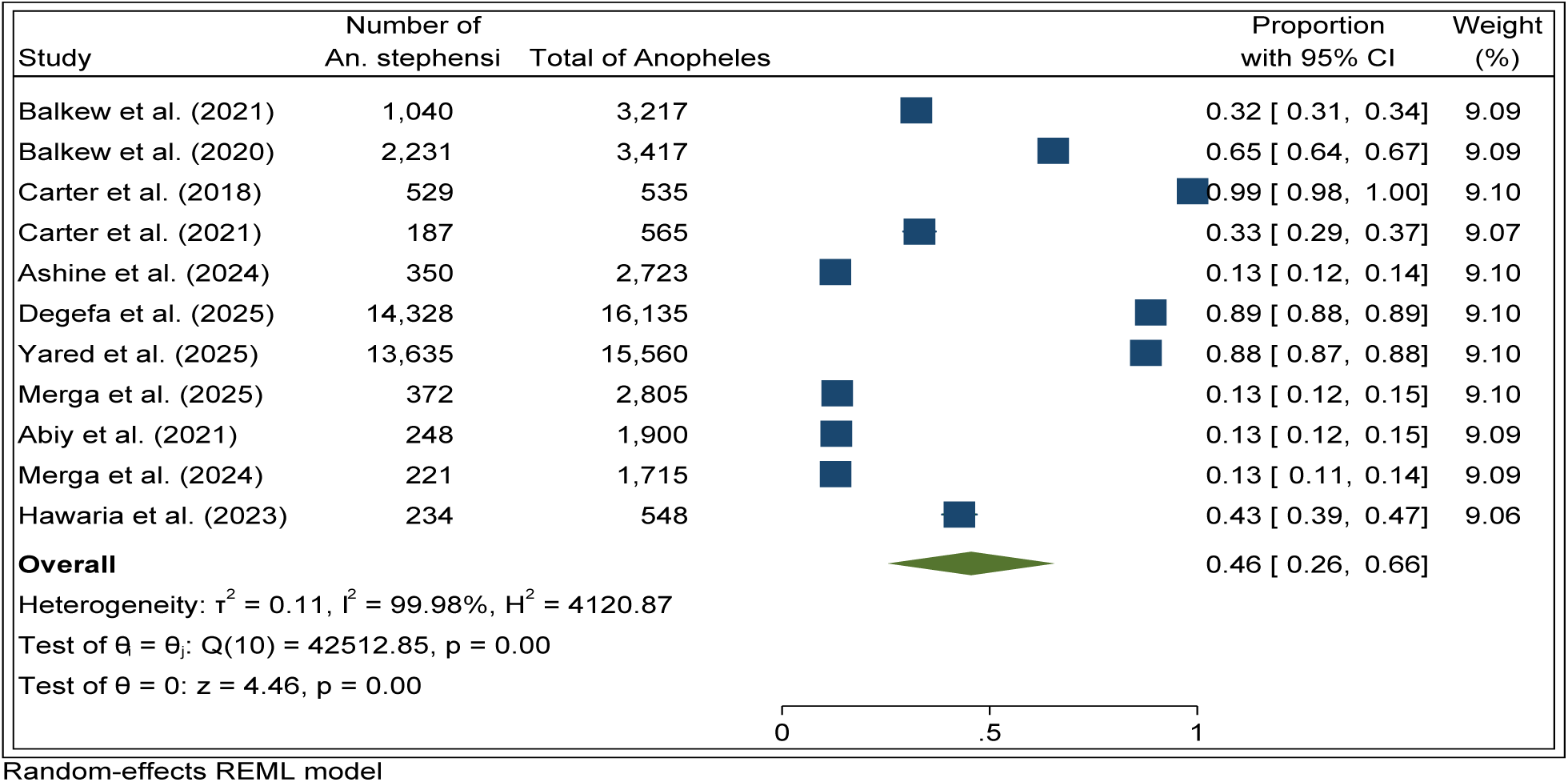
Forest plot of pooled bionomic proportion of *An. stephensi* across ecological and behavioural contexts in Ethiopia

### Subgroup analysis by Ecological Zone and Habitat type of the bionomics studies

Subgroup analysis of bionomics studies revealed significant geographical differences in the proportion of *An. stephensi* among total *Anopheles* mosquitoes. In Eastern Ethiopia, individual studies reported moderate estimates of 0.32 [22] and 0.33 [40], with an overall pooled proportion of 0.60 (95% CI: 0.35–0.85). In Southern Ethiopia, *An. stephensi* was less dominant, with a proportion of 0.43 reported by Hawaria et al. [18]. Studies from Central Ethiopia indicated a lower proportion of approximately 0.13, as reported by Merga et al. [28], Abiy et al. [37], and Ashine et al. [25], with a pooled proportion of 0.13 (95% CI: 0.12–0.14). The overall pooled proportion across ecological zones was 0.46 (95% CI: 0.26–0.66), with significant heterogeneity (τ² = 0.11, I² = 99.98%; Figure 8).

**Figure 8.**
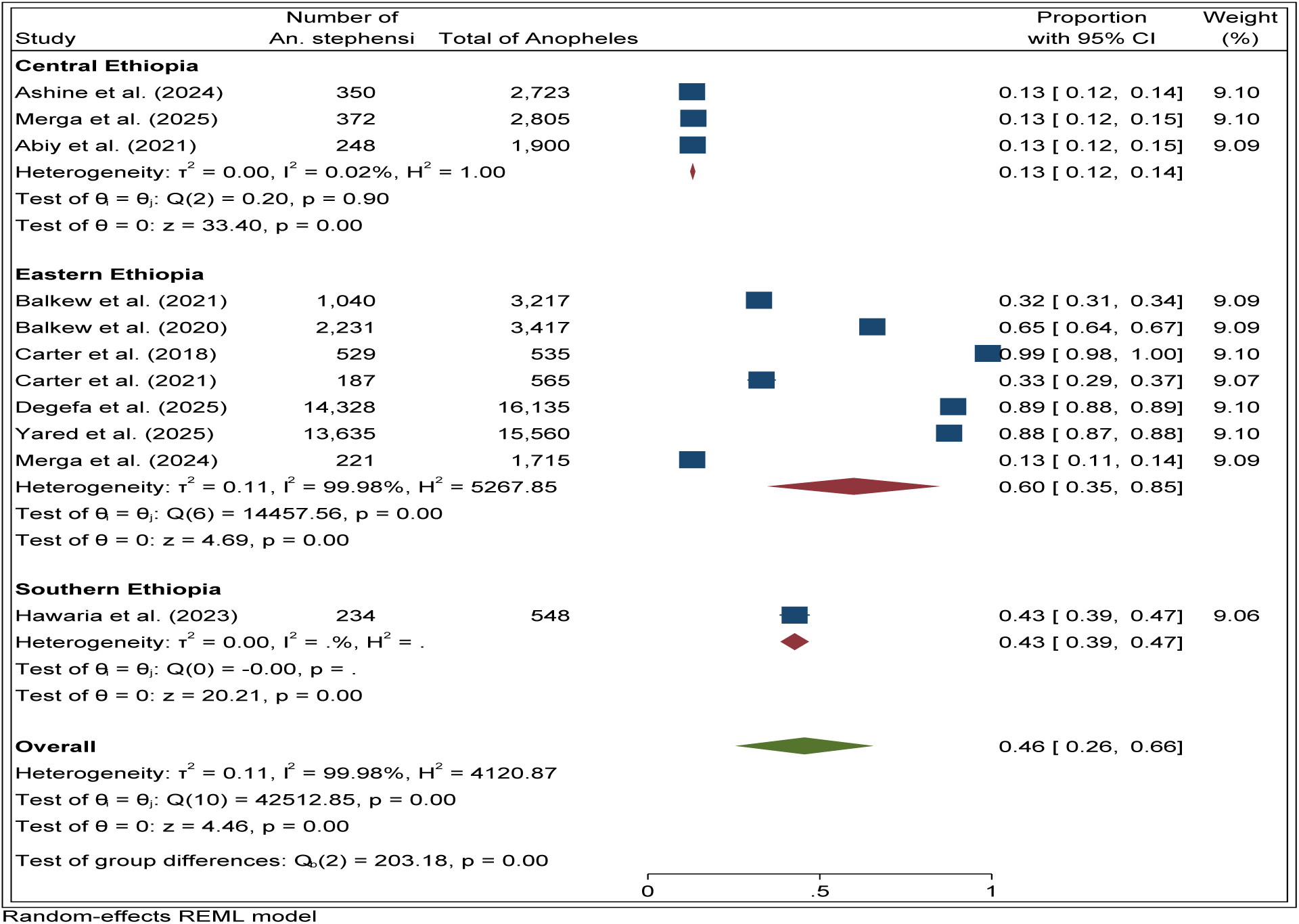
Forest plot of subgroup meta-analysis of *An. stephensi* proportions by ecological zone in Ethiopia

Similarly, subgroup meta-analysis by habitat type showed variation in the proportion of *An. stephensi*. In artificial habitats, pooled estimates ranged from 0.13 [25, 28] to 0.89 [35], with a pooled proportion of 0.45 (95% CI: 0.22–0.67; z = 3.88, p < 0.001). In mixed natural and artificial habitats, estimates varied from 0.13 [37] to 0.99 [33], with a pooled proportion of 0.48 (95% CI: 0.03–0.99; z = 1.85, p = 0.06). Heterogeneity was substantial (τ² = 0.20, I² = 99.98%). The overall pooled proportion across all habitat types was 0.46 (95% CI: 0.26–0.66), with no statistically significant subgroup differences (Q(1) = 0.02, p = 0.90; Figure 9).

**Figure 9.**
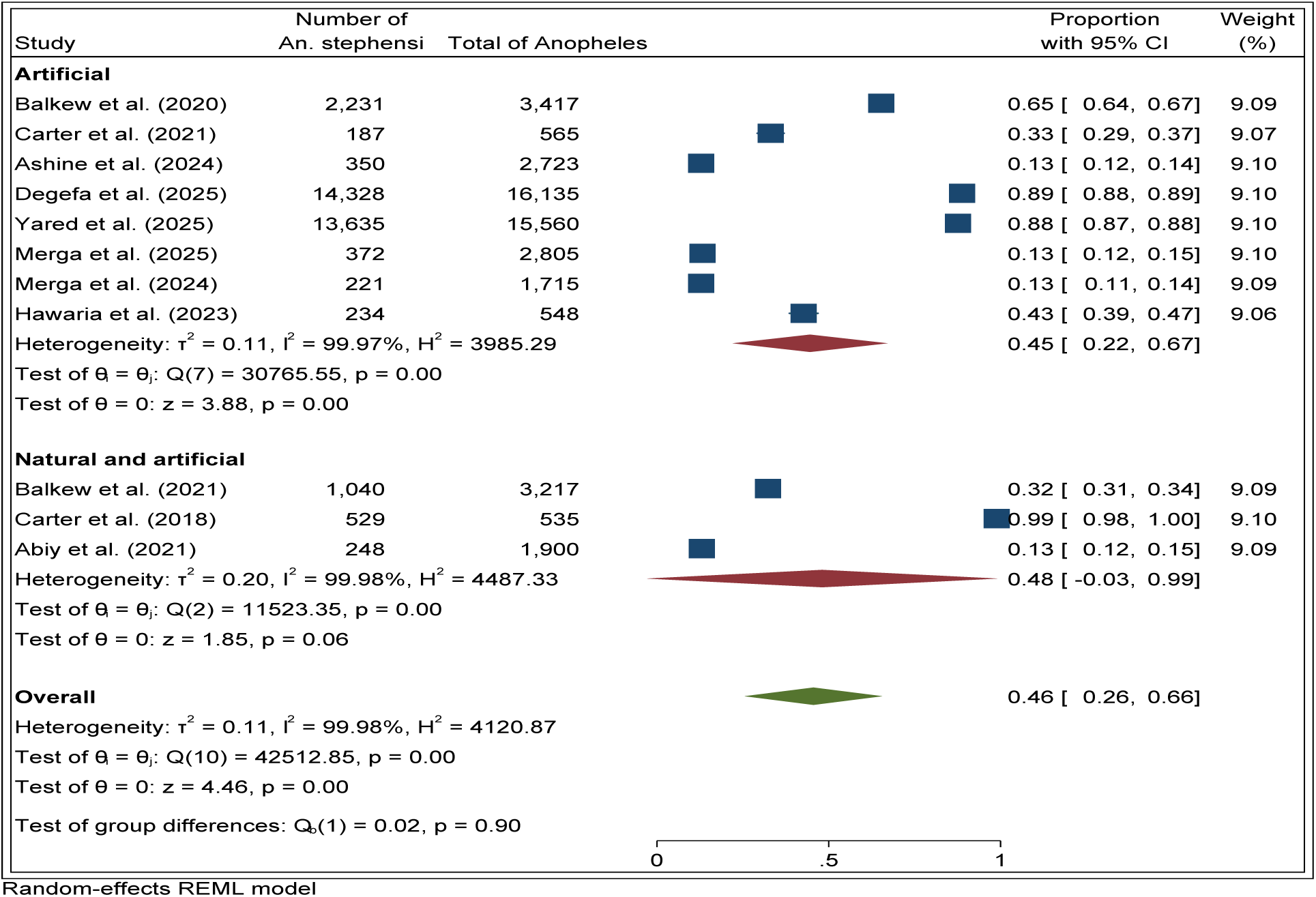
Forest plot of subgroup meta-analysis of *An. stephensi* proportions by habitat type (artificial vs. mixed habitats).

### Subgroup Feeding Preference

Subgroup meta-analysis by feeding preference showed significant variation in the proportion of *An. stephensi*. In the human-associated habitat subgroup, proportions ranged from 0.13 [28, 20] to 0.89 [35], with a pooled estimate of 0.34 (95% CI: 0.07–0.61; z = 2.48, p = 0.01). The variation in *An. stephensi* feeding preference (0.34 for human-associated vs. 0.59 for mixed habitats) and resting behaviour (0.42 for indoor only vs. 0.52 for indoor and outdoor) highlights its behavioural plasticity, which challenges traditional vector control strategies. In contrast to *An. arabiensis*, *An. stephensi* displays opportunistic zoophily and exophily, enabling it to bypass interventions such as treated nets and indoor residual spraying. Heterogeneity was substantial (τ² = 0.11, I² = 99.97%). In the mixed habitat subgroup, proportions ranged from 0.32 [38] to 0.99 [33], with a pooled estimate of 0.59 (95% CI: 0.31–0.87; z = 4.14, p < 0.001). Heterogeneity was high (τ² = 0.10, I² = 99.96%). The overall pooled proportion was 0.46 (95% CI: 0.26–0.66), with no statistically significant subgroup differences (Q(1) = 1.53, p = 0.22; Figure 10).

**Figure 10.**
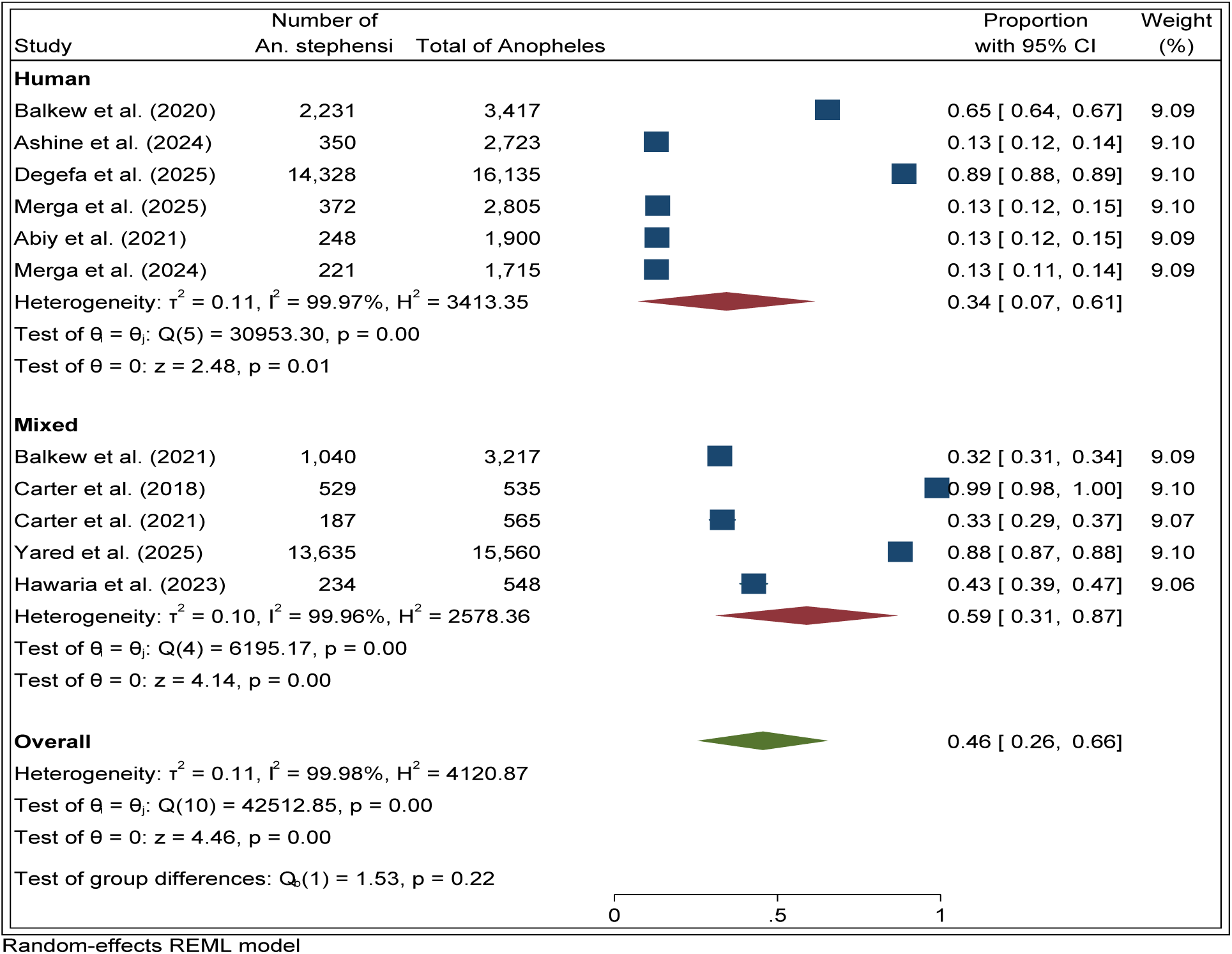
Forest plot of subgroup meta-analysis of *An. stephensi* proportions by feeding preference (human-associated vs. mixed habitats) in Ethiopia

### Subgroup by resting preference

Subgroup meta-analysis by resting preference revealed distinct patterns of the proportion of *An. Stephensi* among total *Anopheles* mosquitoes. Indoor studies reported proportions ranging from 0.13 [25, 20, 37] to 0.89 [2], with a pooled estimate of 0.42 (95% CI: 0.15–0.69; z = 3.01, p < 0.001). Heterogeneity was marked (τ² = 0.14, I² = 99.98%). Indoor and outdoor studies reported proportions from 0.32 [38] to 0.99 [33], with a pooled estimate of 0.52 (95% CI: 0.21–0.83; z = 3.25, p < 0.001). Heterogeneity remained high (τ² = 0.10, I² = 99.90%). The overall pooled proportion across resting preferences was 0.46 (95% CI: 0.26–0.66), with no statistically significant subgroup differences (Q(1) = 0.21, p = 0.64; Figure 11).

**Figure 11.**
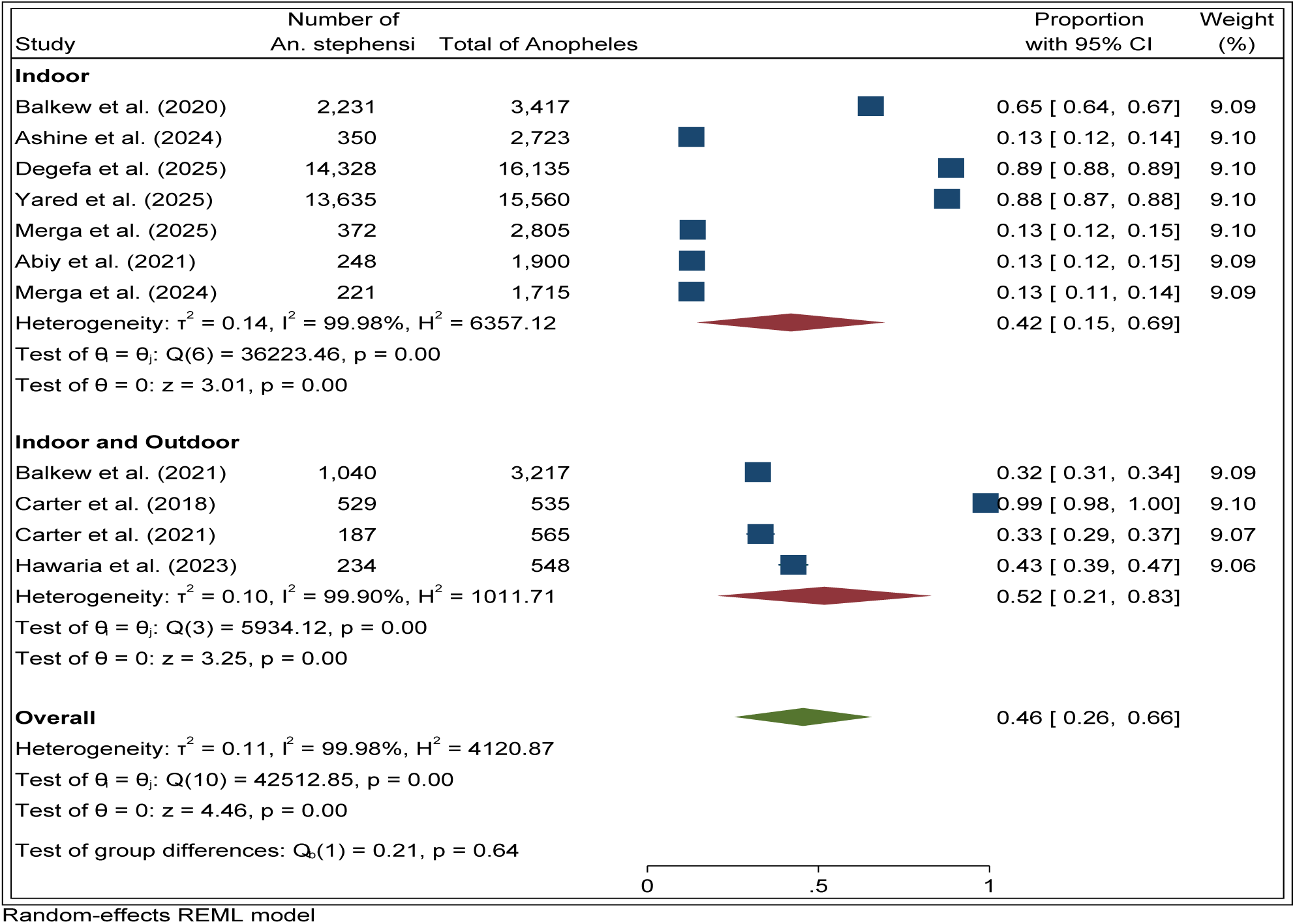
Forest plot of subgroup meta-analysis of *An. stephensi* proportions by resting preference (indoor vs. indoor + outdoor)

### Seasonal design

A subgroup meta-analysis of seasonal collection periods identified significant variations in the proportion of *An. stephensi* mosquitoes among total Anopheles populations. Seasonal studies reported proportions from 0.13 [25, 28, 20] to 0.99 [33], with a pooled estimate of 0.43 (95% CI: 0.16–0.70; z = 3.12, p < 0.001). Heterogeneity was extreme (τ² = 0.13, I² = 99.98%). Year-round studies reported proportions from 0.13 [37] to 0.88 [36], with a pooled estimate of 0.50 (95% CI: 0.17–0.82; z = 3.00, p < 0.001).

Heterogeneity remained high (τ² = 0.11, I² = 99.95%). The overall pooled proportion across seasonal designs was 0.46 (95% CI: 0.26–0.66), with no statistically significant subgroup differences (Q(1) = 0.10, p = 0.76; Figure 12).

**Figure 12.**
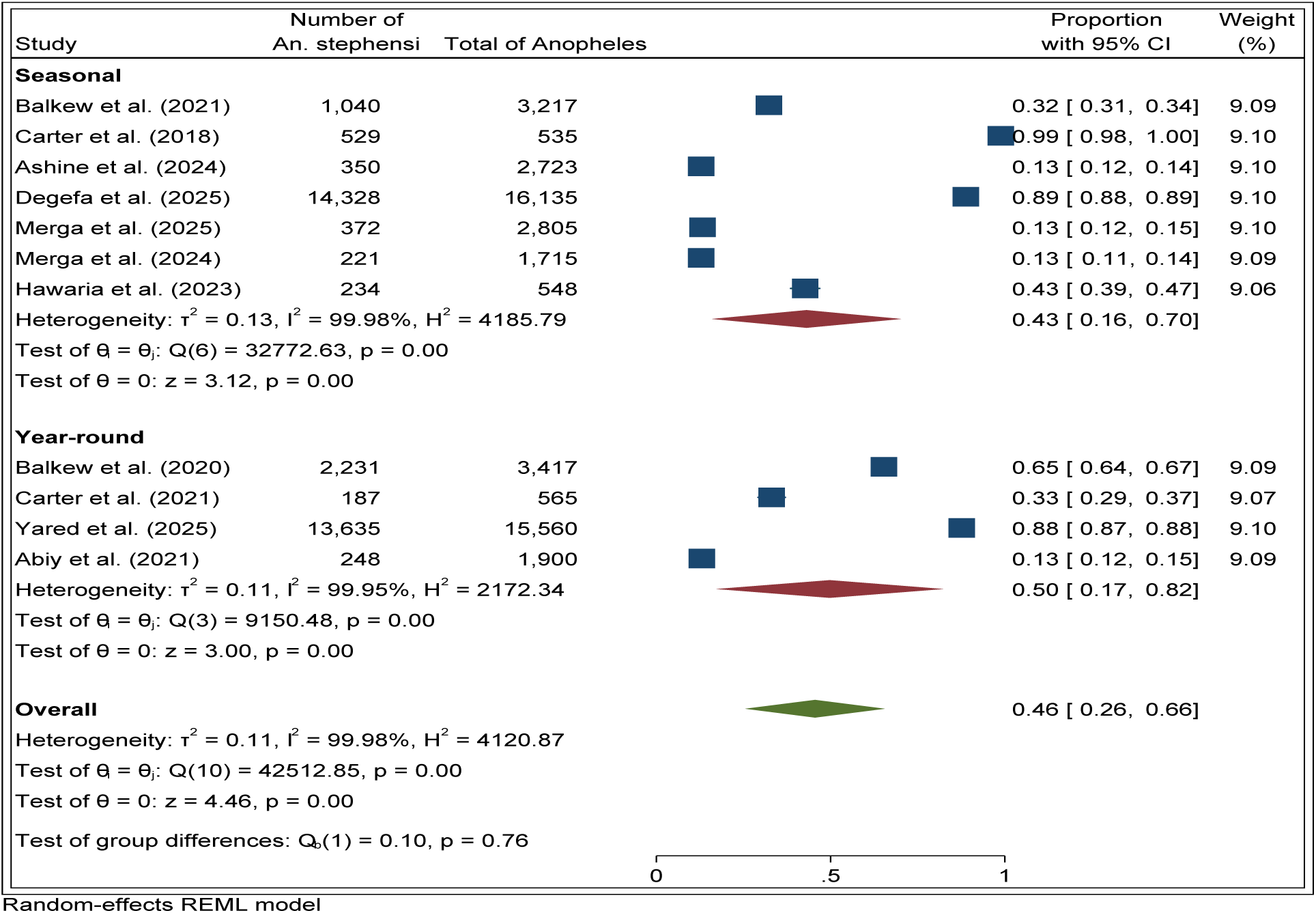
Forest plot of subgroup meta-analysis of *An. stephensi* proportions by seasonal collection design (seasonal vs. year-round)

### Publication Bias and Funnel plot

The funnel plot for the *An. stephensi* bionomics dataset showed no evidence of publication bias, with symmetrical distribution of studies around the pooled estimate (Figure 13). Egger’s regression (β₁ = –18.94, z = –1.11, p = 0.27) and Begg’s test (Kendall’s score = –7.00, z = –0.62, p = 0.64) were non-significant. Trim-and-fill analysis indicated no missing studies, with a stable pooled estimate of 0.455 (95% CI: 0.255–0.655).

**Figure 13.**
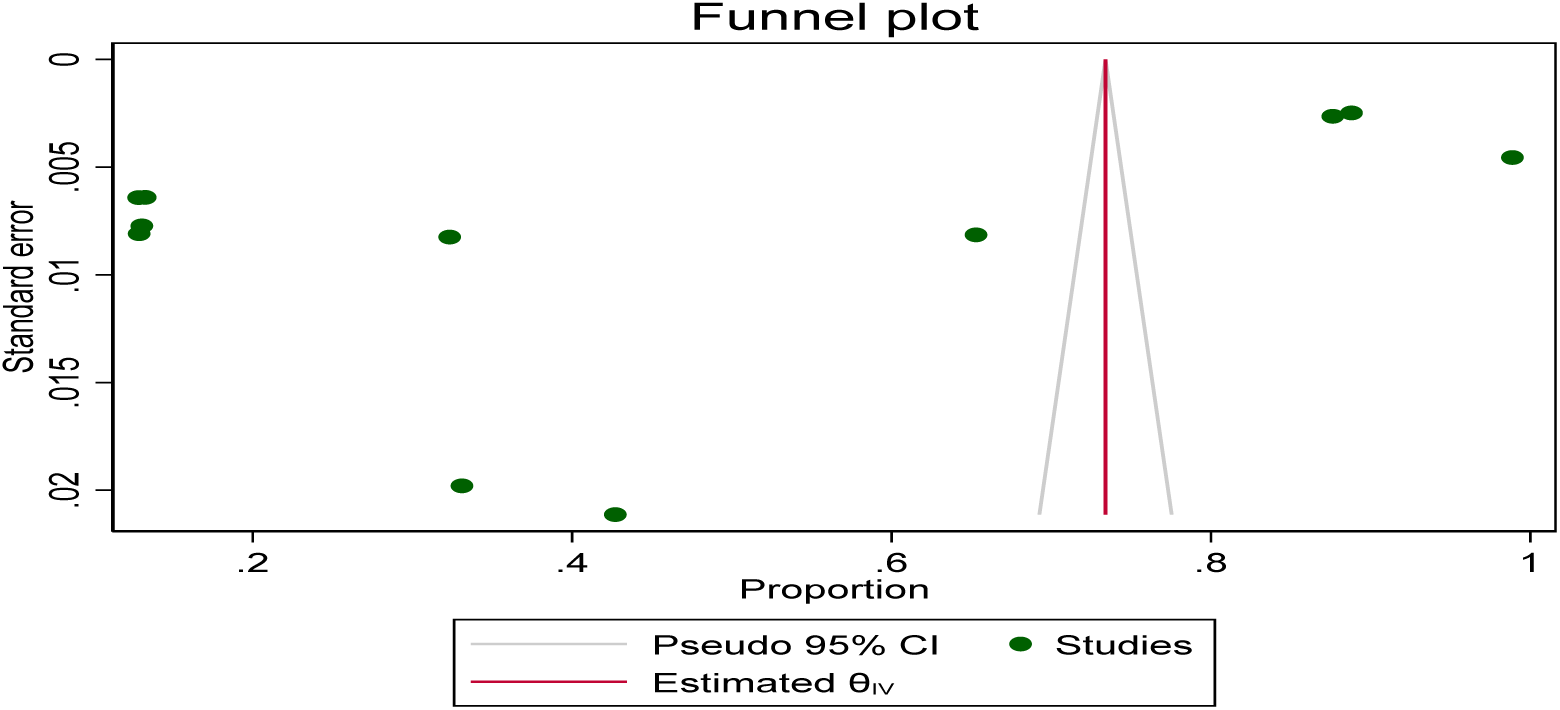
Funnel plot assessing publication bias in bionomic studies of *An. stephensi* in Ethiopia.

## Discussions

The rapid spread of *An. stephensi* in African urban areas poses a significant threat to malaria prevention efforts, potentially reversing decades of progress [49, 50, 51]. *An. stephensi* is rapidly establishing itself in Ethiopian urban areas due to its biological traits and the ecological conditions of urbanisation. It thrives in artificial water sources, exacerbated by poor infrastructure, in stark contrast to *An. arabiensis* and *An. funestus*, which prefer rural areas. *An. stephensi* ability to transmit *P. falciparum* and *P. vivax* increases the susceptibility of urban regions to malaria, while its behavioural traits, including early-evening feeding habits, undermine traditional malaria control methods like LLINs and IRS, resulting in decreased effectiveness [17, 52]. Ethiopia’s invasion of *An. stephensi* illustrates a significant geographic expansion of this species across three continents [52]. Detected in Djibouti in 2012, *An. stephensi* quickly dominated local Anopheles populations, displacing native species and causing malaria outbreaks in areas previously under control [53, 54]. Sudan experienced similar rapid expansion between 2019 and 2021 [55], while in India, stable urban populations have emerged with insecticide resistance and a high vectorial capacity, representing a sustained invasion where *An. stephensi* exceeds 80% of Anopheles populations. In Iran and Saudi Arabia, *An. stephensi* populations exhibit seasonal persistence and notable peridomestic establishment [14].

This systematic review and meta-analysis revealed that *An. stephensi* has quickly emerged as the dominant malaria vector in Ethiopia and the Horn of Africa. The overall pooled epidemiological proportion was 0.51 (95% CI: 0.28-0.75), while the overall pooled bionomics proportion was 0.46 (95% CI: 0.26-0.66). These findings suggest that the species has swiftly evolved in urban and peri-urban habitats, with reported proportions ranging from a low presence of 0.07-0.08 [41, 46] to near-total domination of 1.00 [32]. Emiru et al. [34], Hamlet et al. [31], and Merga et al. [20] all reported high proportions (≥0.8), showing their epidemiological relevance. Moderate proportions (0.27-0.37) in transitional zones [43, 45] indicate coexistence with other Anopheles species.

Geographic heterogeneity implies that south-eastern Ethiopia has nearly complete dominance [32, 43]. On the other hand, Eastern Ethiopia frequently reported high proportion rates when compared to [34, 35, 36], but Central Ethiopia had low proportions of 0.13 [25, 37]. These findings highlight how *An. stephensi* thrives in urbanised ecosystems modified by human activity, notably water storage and peri-domestic habitats showed comparable variability over the Horn of Africa [52].

In Ethiopia, its dominance has increased from 0% in 2016 to 0.51–0.73 by 2024–2025, reflecting a transition from early-stage invasion (Djibouti 2012–2017) to an established invasion similar to that in India today. During the period from 2016 to 2024, Ethiopia’s progression aligns closely with Djibouti’s vector replacement timeline of 5 to 8 years. This correlation indicates that Ethiopia could potentially follow a similar epidemiological trajectory as Djibouti. If the current trend persists, native vector displacement (> 90% *An. stephensi)* may happen by 2028–2032, with potential malaria outbreaks noted in Djibouti, where urban malaria cases surged 10–50 times during the transition. This analysis allows for the prediction of Ethiopia’s future direction and the necessary adjustments in control strategies.

The marked geographic stratification of *An. stephensi* dominance (Central 0.13, Eastern 0.57, South-eastern 0.73) necessitates region-specific vector control approaches rather than a national uniform strategy. Evidence-based recommendations for each region follow: South-Eastern Ethiopia (high dominance, 0.73 pooled proportions), the urgent action is required for integrated urban vector management due to near-total vector replacement. The Eastern Ethiopia (moderate-high dominance, 0.57 pooled proportions); mixed vector populations exhibit ongoing *An. stephensi* expansion, necessitating transitional control strategies. Central Ethiopia (Low Dominance, 0.13 pooled proportions), currently, is lower vector dominance of *An. stephensi*, however, the potential for rapid range expansion exists due to its successful establishment in lower-altitude neighbouring regions.

Ethiopia’s trajectory reflects invasion tendencies in surrounding East African countries and throughout the world. In Djibouti, native vectors were displaced within 5-8 years after identification, with *An. stephensi* accounting for 0.80-0.95 [53, 54]. Similar patterns have been seen in Sudan [55] and Somalia [56]. In India, the species thrives in artificial environments like water storage tanks and building sites, with proportions of around 0.80 [57]. Reports from Iran and Saudi Arabia demonstrate its adaptation to peridomestic habitats and seasonal persistence (14). Climate suitability models forecast increasing spread throughout Africa, exposing millions to urban malaria risk [19]. The bionomics analysis highlights *An. stephensi* ecological adaptability.

The species prefers artificial breeding sites [33, 35], is adaptable to both human-only and mixed-host habitats [40, 42], may be found in both indoor and outdoor resting locations [20, 36], and is most active in the evening and night [34, 45]. Seasonal persistence has been shown in both seasonal and year-round samples [25–37].

This behavioural variability challenges traditional management strategies established for rural vectors like *An. arabiensis* and *An. funestus* [25, 58]. The spread of *An. stephensi* in urban environments creates serious public health concerns. Standard interventions like long-lasting insecticidal nets (LLINs) and indoor residual spraying (IRS) are ineffective due to the mosquito’s exophagic and exophagic behaviour [21, 27]. Rapid urbanisation accelerates its growth, but resistance to pyrethroids and carbamates complicates control efforts. To address these challenges, urgent policy action is essential. Priority actions include improving urban entomological surveillance to monitor *An. stephensi* populations, managing artificial breeding sites through larval source management, implementing integrated vector management with environmental modification, biological control, and insecticide use, and promoting community-based interventions to reduce breeding habitats. Integrated urban vector management in Ethiopia, combining LLINs, larval source management, insecticide resistance monitoring, and community engagement, faces significant implementation barriers within the health system.

This systematic review identifies critical research gaps in vector control optimisation related to *An. stephensi* in Ethiopia. Key priorities include: (1) Geographic Coverage Gaps; limited studies in central and south-western Ethiopia necessitate standardised entomological surveys across districts; (2) Temporal Dynamics—there’s a lack of longitudinal studies on *An. stephensi*; establishing 5-year cohort studies in urban sites is essential to track population dynamics and invasion patterns; (3) Clinical Malaria Attribution—absence of studies quantifying malaria linked specifically to *An. stephensi* hinders health impact assessments; case-control studies are needed to estimate its burden; (4) Intervention Trials no randomised controlled trials (RCTs) have evaluated urban vector management; multi-site RCTs comparing various interventions are required; (5) Insecticide Resistance limited resistance mechanism data underscore the need for whole-genome sequencing to inform resistance management strategies; (6) Urban Ecology inadequate data on microhabitat preferences call for standardised surveys in urban districts to optimise larval control; (7) Behavioural Ecology limited quantitative data necessitate detailed studies on feeding and resting behaviours to enhance protection strategies

This systematic review and meta-analysis demonstrates notable methodological strengths, including compliance with PRISMA 2020 guidelines and PROSPERO registration, which enhance transparency and reduce selective reporting. It provides comprehensive vector characterisation through the dual reporting of epidemiological proportions and bionomic characteristics, alongside rigorous study quality appraisal. The evaluation of publication bias revealed no significant issues; however, extreme heterogeneity (I² = 99.98%) limits the generalizability of the pooled results, reflecting ecological diversity and methodological variability. Detection bias in studies of *An. stephensi* arises from their focus on regions where the species is suspected or previously detected, leading to potential overestimation of vector proportions due to systematic under-sampling in areas with low or absent populations.

Additionally, temporal confounding in these studies, conducted between 2016 and 2025, cannot distinctly attribute geographic variance in vector proportions to ecological heterogeneity or recent invasions, necessitating assumptions about ecological suitability. The heterogeneity in surveillance methods ranging from larval to adult collection and indoor versus outdoor sampling further compounds the variability and results in high heterogeneity estimates. Furthermore, while vector presence is documented, there is a lack of clinical outcome data linking *An. stephensi* to malaria incidence or parasitemia burden, typically relying on proportional assumptions that may not accurately reflect risk.

## Conclusion

This meta-analysis demonstrates that *An. stephensi* is widely distributed throughout Ethiopia and the Horn of Africa, with pooled proportions indicating high proportions across epidemiological and ecological settings. Its adaptability to various ecological zones, habitats, host preferences, resting areas, feeding periods, and seasons demonstrates its invasive ability. Ethiopia’s trajectory is consistent with worldwide invasion patterns, especially in Djibouti and India, where *An. stephensi* has shifted malaria transmission dynamics. Integrated monitoring and customised vector control strategies, focusing on artificial breeding sites and urban areas, are urgently needed. Without immediate action, *An. stephensi* threatens to undercut Ethiopia’s malaria eradication efforts and increase urban malaria spread throughout Africa.

## Abbreviationsw

CI: Confidence Interval
CRD: Centre for Reviews and Dissemination (used in PROSPERO registration number)
IRS: Indoor Residual Spraying
I²: I-squared statistic (measure of heterogeneity)
JBI: Joanna Briggs Institute
LLINs: Long-Lasting Insecticidal Nets
N: Total number of *Anopheles* collected
n: Number of *An. stephensi* collected
PRISMA: Preferred Reporting Items for Systematic Reviews and Meta-Analyses
PRISMA-P: Preferred Reporting Items for Systematic Reviews and Meta-Analyses Protocol
PROSPERO: International Prospective Register of Systematic Reviews
Q statistic: Test for subgroup differences in meta-analysis
τ² (Tau-squared): Between study variance in meta-analysis
θ_IV: Inverse-variance weighted estimate
WHO: World Health Organization
Z: z-score (test statistic in meta-analysis)

## Acknowledgements

We want to thank our colleagues who had a great contribution to the preparation of this manuscript.

## Authors’ Contributions

T.B. directed the systematic review and meta-analysis, designed the study, selected the articles, developed the protocol, registered it with PROSPERO, extracted the data, performed the statistical analysis, and prepared the manuscript. Database searches and study selection were conducted by T.B., K.L., and A.D. Data extraction and quality appraisal using JBI tools, as well as statistical analysis and meta-analysis in STATA, were performed by T.B., K.L., A.D., and L.R. The initial manuscript draft was prepared by T.B., while K.L. and A.D. provided critical revisions. All authors read and approved the final version of the manuscript.

## Data availability

The data that support the findings of this study are found in the manuscript.

## Declarations

### Ethics approval and consent to participate

Not applicable.

### Consent for publication

Not applicable.

### Clinical trial

Not applicable.

